# The fluctuation-based regime of thalamocortical circuitry

**DOI:** 10.1101/2025.11.20.689511

**Authors:** Liam Jennings, David Hansel, Nicholas J. Priebe

**Affiliations:** The University of Texas at Austin; Cerebral Dynamics, Plasticity and Learning Laboratory, UMR8002, CNRS, Univ. Paris Cité

## Abstract

Neocortical neurons in primary visual cortex (V1) integrate excitatory inputs from both thalamic and cortical sources to generate selectivity for orientation, direction and disparity. Feedforward models have posited that thalamocortical input can drive these selectivities, but the relative sparsity of thalamic synapses raises questions about the sufficiency of this input. To determine whether thalamic drive is sufficient to evoke cortical responses, we quantified the current threshold required to evoke action potentials in vivo using whole-cell recordings in mouse V1. We then isolated thalamic input via optogenetic cortical silencing and compared its strength to the measured spiking threshold. We find that although average thalamic input is below the threshold for action potential generation, trial-by-trial fluctuations in thalamic drive are sufficient to evoke visually driven action potentials. These results demonstrate that thalamic excitation alone can drive cortical spiking, but its effectiveness depends on fluctuations in synaptic input.

## Introduction

Neocortical neurons receive thousands of excitatory synaptic inputs. Among those, 90% are from other cortical neurons and 10% from thalamocortical projections. (Peters & Payne, 1993; Douglas et al. 1995; Binzegger et al., 2004). In primary visual cortex (V1), these thalamic and cortical inputs are integrated to generate selectivity for diverse features such as orientation, direction, and binocular disparity (Hubel & Wiesel, 1962; Priebe & Ferster, 2008). Feed-forward models have been successful in explaining how thalamocortical inputs contribute to the emergence of these feature selectivities (Hubel & Wiesel, 1962; Ferster et al., 1996; Priebe & Ferster, 2005). Many of these models rely on spike threshold non-linearities to shape tuning properties (Priebe & Ferster, 2005; Finn et al., 2007; Priebe & Ferster, 2008; Priebe & Ferster, 2012). Since there is much less thalamocortical than intracortical connectivity, however, it is unclear whether the amplitude of feedforward drive is sufficient to account for cortical selectivity; it is not known if the mean thalamocortical drive alone is sufficient to generate visually-evoked action potentials. Understanding the strength of thalamocortical drive places important constraints on the degree that feedforward drive could shape cortical selectivity.

Given the large disparity in the number of thalamocortical and intracortical synaptic inputs, many models have relied on strong excitatory recurrent connections within cortex to shape selectivity (Ben-Yishai et al., 1995; Douglas et al., 1995). These recurrent models have also provided an account for a number of features in V1, such as direction selectivity, orientation selectivity, and contrast invariance of tuning (Somers et al., 1995; Ringach et al., 1997; Sompolinsky & Shapley, 1997; Somers et al., 1998). In these models the thalamus provides broadly selective input to V1, while recurrent circuitry amplifies and sharpens selectivity. Several of these models rely on strong excitation balanced by strong recurrent inhibition (Hansel & van Vreesewijk, 2012; Mariño et al., 2005; Murphy & Miller, 2009; Pattadkal et al., 2018). Furthermore, cortical inhibition is likely to play a key role in sculpting selectivity and controlling gain (Carandini et al., 1997; Isaacson & Scanziani, 2011; Ahmadian et al., 2013). Indeed, experimental studies have also found that inhibition shapes selectivity (Liu et al., 2011; Tan et al., 2011 Li et al., 2013) particularly in awake animals (Haider et al., 2013), further complicating the role of thalamocortical inputs in generating selectivity in V1.

Since a critical assumption of feedforward models is that the thalamocortical drive is sufficient to evoke responses in the cortex, we set out to determine the strength of thalamic input compared to the drive necessary to evoke action potentials. Importantly, feedforward models have relied on feedforward activity in which the mean thalamic drive is suprathreshold, but no measurement has assessed this assumption (Hubel & Wiesel, 1962, Ferster et al., 1996, Finn et al., 2007). We approach this problem by first defining the spiking threshold of cortical neurons on the basis of how much current is required to increase spike rate *in vivo*. Using whole-cell recordings from mouse V1 we measured the relationship between injected current of different amplitude and duration and spike rate under spontaneous conditions. We systematically extracted the current threshold to elicit action potentials of individual neurons *in vivo*, and show its dependency on the current injection duration and ongoing membrane potential fluctuations. We then performed voltage-clamp recordings to assay the strength of visually evoked synaptic drive onto individual neurons. Through optogenetic inactivation of the cortex, we isolated the thalamic contribution to the total excitatory input as well as its relationship to the current threshold. While the average thalamic drive to cortical neurons is generally smaller than the current threshold of cortical neurons, the trial-by-trial fluctuations in thalamic drive are large, and sufficient to bring V1 neurons to the spiking threshold. Therefore thalamic excitation is sufficient to drive visually driven action potentials in V1, but these action potentials rely on input fluctuations.

## Results

### Measurement of current thresholds *in vivo*

To examine the relationship between current injection and spike rate, we injected current pulses of varying amplitude (0-800pA) during somatic whole-cell recordings in mouse V1 (Figures 1A and 1B). As the integration of synaptic input varies across timescales (Softky, 1994; Shadlen & Newsome, 1998; Stuart et al., 2016), we injected current pulses of two different durations, 20ms and 100ms. The short duration is close to the timescale of the cell membrane, while the longer pulses reflects drive over timescales closer to those elicited by visual stimulation.

**Figure 1:**
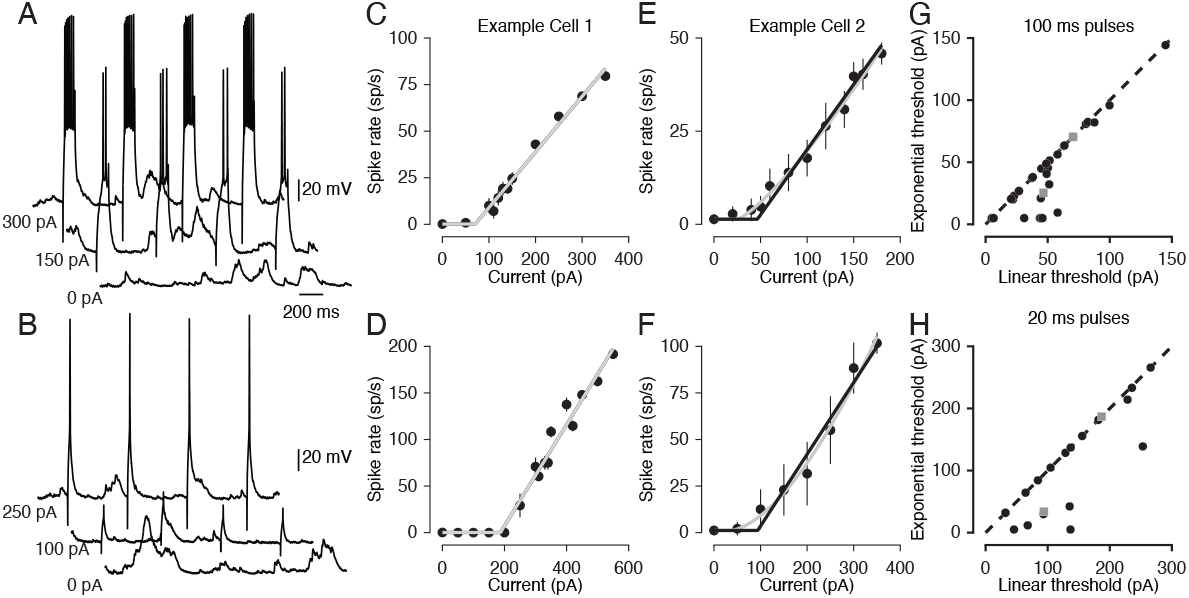
Current thresholds in response to varying current duration can be reliably estimated *in vivo*. (A) Example whole-cell traces for three different current amplitudes for 100ms pulse duration. The cell is the same in the three traces. (B) Same example cell shown in (A), but for the 20ms pulse duration. (C) Frequency-vs-current curve for the cells shown in (A) and (B). Current-pulse duration is 100ms. Solid line: fit to a threshold-linear f-I curve. Error bars are standard errors. (D) Same as in (C), but for 20ms pulses. (E) An example frequency-versus-current curve for a second example cell in response to 100ms current pulses. The exponent, *p*, is a free parameter of the fit constrained by *p >=* 1. (F) Same as in (E), but for 20ms pulses. (G) Comparison of the current threshold for threshold-linear (p=1) and threshold-power law fits (p>1). The duration of the current pulse is 100ms. Gray squares indicate example cells shown in C-F.(H) Same as in (A), but for 20ms pulses.

The spike rate of neurons *in vivo* followed a similar pattern to those measured *in vitro*: for small currents no change in spike rate was observed. Once current amplitude reached a threshold, firing rate increased and saturated for large injection amplitudes (Figures 1C, 1D, 1E, and 1F). To estimate this threshold current, we employed two methods. First, we looked for a statistical difference between the distributions of spikes evoked by the current injection and the distribution observed without current injection. We assayed statistical significance for all injection amplitudes, using Bonferroni correction, and defined the lowest current amplitude to increase rate significantly as the ‘statistical threshold’ current. Second, given that the relationship between injected current and spike rate can be approximated by a rectified power law (Hansel & van Vreeswijk 2002; Miller & Troyer, 2002; Priebe & Ferster, 2008), we fit our data to

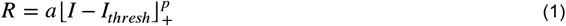

where *R* is the evoked spike rate, *+* is rectification, *p* is the exponent, *I*_*thresh*_ is the fitted threshold current and *a* is a gain factor. We started by fixing the exponent, *p*, to 1 (threshold linear fits, Figures 1C and 1D), which allows for a direct comparison to the statistical threshold. Both of these methods resulted in comparable estimates of threshold current, though the statistical threshold was marginally larger than the fitted one for many of the neurons in our dataset (r = 0.87, p = 2.361e-17, n = 54, Pearson’s correlation).

In response to 100 ms current pulses we found that the threshold current varied across cells between 5 and 150 pA with an average near 50 pA (Table 1, statistical *I*_*thresh*_ = 54.41 ± 27.43, fit *I*_*thresh*_ = 48.96 ± 28.69 pA). The average threshold was significantly higher for 20 ms pulse duration (Figures 2A and 2B; Table 1). In addition, the threshold current was much more variable across the dataset, ranging from 40-350pA (Figures 2A and 2B; Table 1). This may be due to the influence of spontaneous voltage fluctuations resulting from background activity during current injections. Across the sample population in which the threshold was measured for both 100ms and 20ms, it was on average larger by 75.88pA ± 11.73 pA (mean increase for fitted threshold, 70.36 ± 12.43 pA, n = 17; Figure 2C) for 20 ms than for 100 ms. Additionally, the difference in the current threshold between the two durations was statistically significant (statistical I_thresh_: p = 7.731e-06, paired t-test; fit I_thresh_: p = 3.539e-05, paired t-test). These results show that, *in vivo*, the spiking thresholds for single neurons significantly depend on the timescale of the inputs.

**Table 1:** Current threshold statistics.

|  | Overall |  | UP |  | DOWN |  | UP vs DOWN |
| --- | --- | --- | --- | --- | --- | --- | --- |
| | Mean ± SD (pA) | Range (pA) | Mean ± SD (pA) | Range (pA) | Mean ± SD (pA) | Range (pA) | Paired t-test |
| 100ms Fitted | 48.96 ± 28.69 | 5–144.91 | 34.62 ± 27.20 | 5–109.13 | 72.72 ± 37.19 | 5–200 | 1.2276e – 07 |
| 100ms Statistical | 54.41 ± 27.43 | 20–150 | 56.32 ± 25.42 | 20–150 | 80.44 ± 46.54 | 20–250 | 1.4574e – 04 |
| 20ms Fitted | 133.13 ± 72.89 | 31.85–265.82 | 93.11 ± 70.74 | 5–227.49 | 190.49 ± 72.67 | 48.64–300 | 8.4629e – 06 |
| 20ms Statistical | 145.5 ± 83.25 | 40–350 | 136.56 ± 74.10 | 40–280 | 218.13 ± 94.10 | 60–450 | 9.2806e – 04 |
| All Fitted | 80.13 ± 64.03 | 5–265.82 | 55.43 ± 57.88 | 5–250.05 | 117.79 ± 78.80 | 5–300 | |
| All Statistical | 88.15 ± 70.18 | 20–350 | 86.67 ± 68.09 | 20–350 | 132.84 ± 108.44 | 20–550 | |

**Figure 2:**
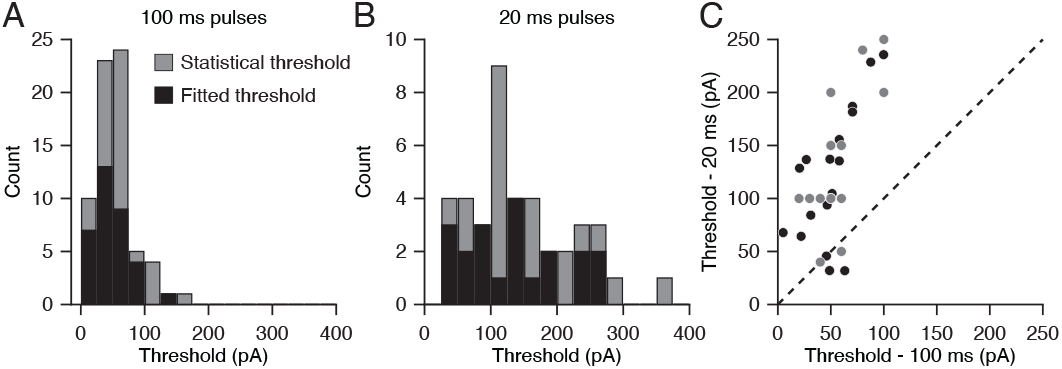
Threshold measurements significantly depend upon the timescale of the injected current. (A) Distribution of fitted and statistical current threshold measurements for the 100ms pulse duration. (B) Same as in (A), but for 20ms. (C) Scatterplot comparing current threshold measurements made with 100ms and 20ms pulse durations in the same neurons.

We next allowed the exponent, *p*, to vary, and extracted the threshold current. For many neurons (22/34, 100ms; 12/20, 20ms) I_thresh_ did not shift (Figures 1G and 1H), but for other cells (12/34, 100ms; 8/20, 20ms) there was a clear reduction in the threshold current. Overall, using an exponent greater than 1 reduced the mean fit threshold from 48.96 to 40.54 pA (17%) and from 133 to 104 pA (22%) for 100 ms and 20 ms pulses, respectively. These reductions in the fit I_thresh_ reduced the correlation between the statistical threshold and the fit threshold, but they were nonetheless strongly related (r = 0.48, p = 0.004, n = 34, for 100ms; r = 0.80, p = 2.492e-05, n = 20, for 20 ms). Given the presence of ongoing noise, a power-law model provides a better fit to the data (Hansel & van Vreeswijk, 2002; Miller & Troyer, 2002), but we will primarily use the threshold linear model (p=1) as it provides a better match to the statistical threshold.

An alternative model for action potential initiation suggests that rather than a constant current strength a constant charge must be applied to generate an action potential (Hodgkin & Rushton, 1946). The difference in threshold between the 20ms and 100ms pulse duration could arise from the difference in charge between conditions. If a charge threshold exists, it should be the same across the measurements for both pulse durations in each cell. The relationship between charge, current, and time is

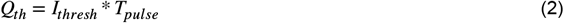

where *Q*_*th*_ is the charge threshold in picocoulombs (pC), *I*_*thresh*_ is the measured threshold in pico-amperes (pA), and *T*_*pulse*_ is the pulse duration in seconds.

For cells in which the threshold was estimated for both 100 ms and 20 ms duration we found that the charge was significantly greater for 100 ms pulses (median charge for 100 ms = 5 pC; median charge for 20ms = 2 pC; p = 7.925e-07, paired t-test, n = 17). As the amount of charge required to generate action potentials differed across pulse durations, neurons in V1 do not appear to have a charge threshold; differences in current threshold between the two pulse durations are likely due to other factors, such as the intrinsic properties of the neurons and network activity.

Intrinsic properties play a key role in shaping the input-output relationship of neurons *in vivo* (Li et al., 2020). In particular, membrane depolarization in response to stimulus is largely governed by input resistance (Hodgkin & Huxley, 1952; Troyer & Miller, 1997). We examined whether the threshold current was related to input resistance (R_n_) across our sample population. To determine input resistance, we measured the subthreshold membrane potential responses to 100ms current injections of 100 pA. These responses were then fit to a double exponential (Anderson et al. 2000a)

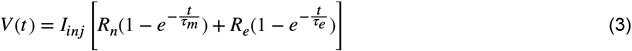

where the first term accounts for the cell’s membrane integration and the second one for the faster effect of the electrode. Across our entire dataset, the mean input resistance was 177 ± 10 MΩ and the mean membrane time constant (***τ***_m_) was 21 ± 0.63 ms. For both 100ms and 20ms, R_n_ was negatively correlated with current thresholds (100ms statistical significance threshold: r = -0.30, p = 0.0868, 100ms fitted threshold: r = -0.36, p = 0.0371, n = 34, Pearson’s correlation; 20ms statistical significance threshold: r = -0.40, p = 0.1073, 20ms fitted threshold: r = - 0.55, p = 0.0218, n = 17, Pearson’s correlation; Figures 3A and 3B). Similar to measurements of R_n_, we see a negative correlation between ***τ***_m_ and thresholds (100ms statistical I_thresh_: r = -0.33, p = 0.0541, 100ms fit I_thresh_: r = -0.51, p = 0.002, n = 34; 20ms statistical I_thresh_: r = -0.32, p = 0.2095, 20ms fit I_thresh_: r = -0.43, p = 0.088, Pearson’s correlation; Figures 3C and 3D).

**Figure 3:**
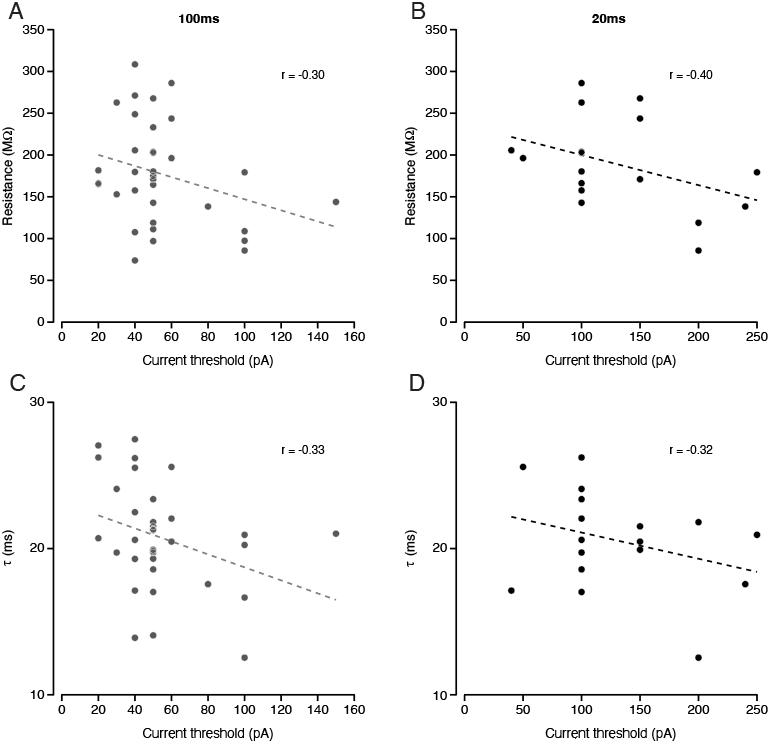
Measured threshold and intrinsic properties are weakly inversely related. (A)The relationship between input resistance and measured statistical thresholds for the 100ms pulse duration. (B) Same as in (A), but for the 20ms pulse duration. (C) Same as in (A), but for neuronal time constant. (D) Same as in (C) for the 20ms pulse duration.

We provide a descriptive metric of current threshold for visual cortical neurons *in vivo* and show that varying the duration of current injection has a significant impact on measured thresholds. Furthermore there is a weak relationship between intrinsic properties (R_n_ and ***τ***_m_) and the current threshold.

### The strength of thalamic drive relative to the threshold

Prior studies have indicated that thalamic input to V1 neurons is relatively weak and therefore requires intracortical excitatory amplification (Douglas et al., 1995; Stratford et al., 1996). While recurrent connections in V1 can allow for amplification of feedforward signals, a fundamental question regarding the strength of feedforward input remains: can thalamic feedforward input alone evoke spiking activity within V1?

Given that the current threshold across neurons varied, we quantified the amount of direct input each of these neurons receives from the thalamus during visual stimulation at each neuron’s preferred orientation. To isolate excitatory inward currents, we held neurons near the reversal potential for inhibition (-75mV to -80 mV) in voltage clamp. Recordings were made in animals in which ChR2 was expressed in PV+ neurons, allowing us to inactivate the cortical circuit during stimulation using light (Lien & Scanziani, 2013). Invariably the visually-evoked responses of neurons were weaker during cortical inactivation (Figure 4A). To estimate the change in the response amplitude we first cycle-averaged the current traces at the temporal frequency of the visual stimulus (Figure 4B). We then computed the peak response amplitude, defined as the sum of the mean and first Fourier coefficient of the response.

**Figure 4:**
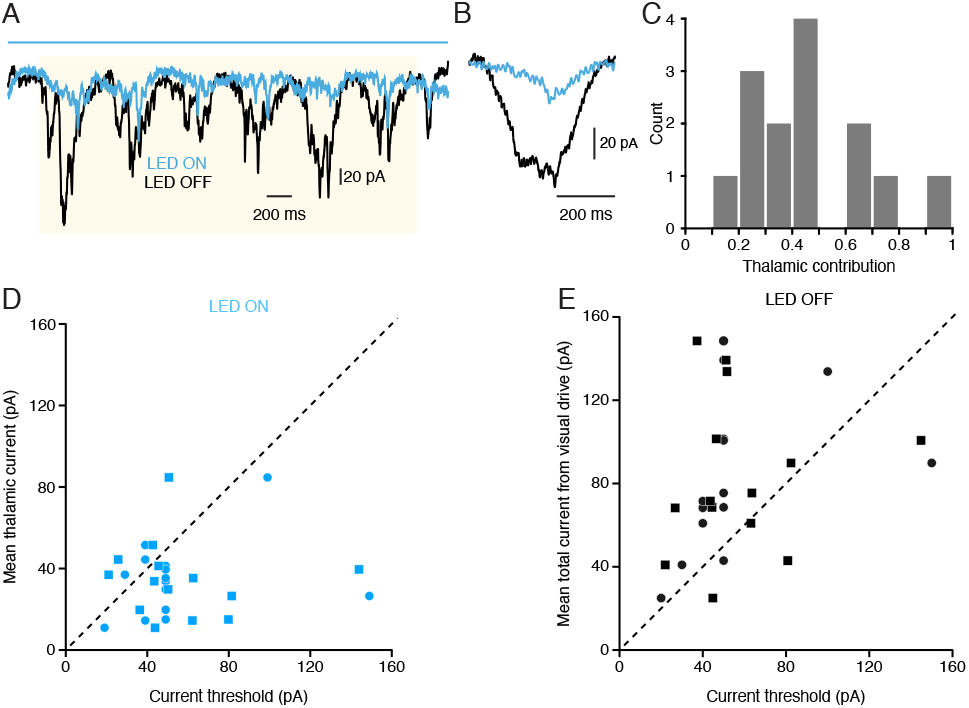
Relationship between feedforward thalamic drive and measured thresholds. (A) Example voltage-clamp trace in response to grating stimulation at the optimal orientation during control conditions (black) and optogenetic inactivation of cortex (blue). (B) Example cycle-averaged response for the cell shown in (A) during control (black) and optogenetic inactivation of cortex (blue). (C) Histogram of the distribution of measured thalamic contribution to visually-evoked activity each neuron receives. (D) Scatterplot comparing the measured thalamic current during cortical inactivation to the measured statistical rheobase (filled circles) and fitted rheobase (open squares). (E) Same as in D, but for the control.

To determine whether the thalamic drive alone is sufficient to generate action potentials in response to a drifting grating stimulus we compared the peak input amplitude to the current statistical and fit thresholds. Because the presented drifting grating stimulus was slow (2 Hz) we used the thresholds measured for the 100 ms stimulus (Figure 4A). For those neurons that received direct thalamic input, it comprised 40% ± 6% of the total current measured when the cortex was intact (Fig. 4C). The mean peak amplitude of the cycle-averaged thalamic drive was subthreshold for the majority of neurons (11/14 neurons for statistical I_thresh_ ; 10/14 neurons for fit I_thresh_ ; Figure 4D). Across cells, peak thalamic drive comprised ∼60% of the current required to reach action potential threshold. In comparison, the total visually evoked current input, measured when the cortex was intact, was supra-threshold for the majority of neurons (n = 12/14 cells for statistical I_thresh_; n = 12/14 cells for fit I_thresh_, Figure 4E; Table 2). Importantly, the thalamus contributed less than half to the total input.

**Table 2:**
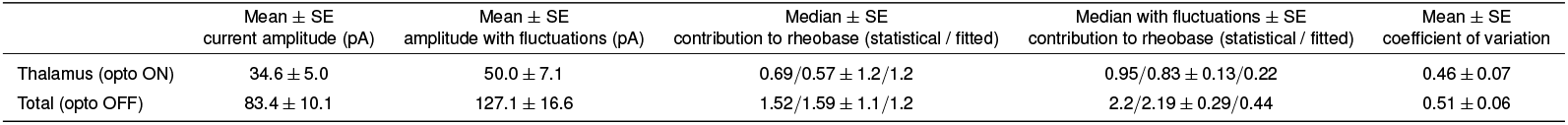
Cortical inactivation statistics.

| | Mean ± SE<br>current amplitude (pA) | Mean ± SE<br>amplitude with fluctuations (pA) | Median ± SE<br>contribution to rheobase (statistical / fitted) | Median with fluctuations ± SE<br>contribution to rheobase (statistical / fitted) | Mean ± SE<br>coefficient of variation |
| --- | --- | --- | --- | --- | --- |
| Thalamus (opto ON) | 34.6 ± 5.0 | 50.0 ± 7.1 | 0.69/0.57 ± 1.2/1.2 | 0.95/0.83 ± 0.13/0.22 | 0.46 ± 0.07 |
| Total (opto OFF) | 83.4 ± 10.1 | 127.1 ± 16.6 | 1.52/1.59 ± 1.1/1.2 | 2.2/2.19 ± 0.29/0.44 | 0.51 ± 0.06 |

Our measurements indicate that the mean thalamic drive is not sufficient to reach the current threshold for spiking. The thalamic drive fluctuated substantially on a cycle-to-cycle basis, however, and these fluctuations may be a key component for neurons to reach threshold. For many neurons, while the mean response remained below threshold, the cycle-to-cycle fluctuations resulted in current responses above threshold (Figure 5A ; Table 2). To better understand the relationship between cycle-to-cycle fluctuations and current threshold, we measured the standard deviation of the cycle average thalamic input at each time point relative to the start of the cycle. The summation of the mean and standard deviation was often about the current threshold (Fig. 5A). We then summed the mean and modulation amplitude for each stimulus cycle and measured the amplitude at the time of the peak response (Finn et al., 2007). The inclusion of these feedforward fluctuations pushes thalamic input above the current threshold for many of the cells in our dataset (n = 7/14 cells for statistical I_thresh_, n = 6/14 cells for fit I_thresh_, Figures 5B and 5C; Table 2). The thalamic drive averaged across all recorded neurons was 34.6 pA ± 5.0 pA. When taking into account fluctuations, it increased to 50.0 pA ± 7.1 pA. We also observed this increase in mean drive across our sample population for measurements of the total current during visual stimulation (without fluctuations: 83.4 pA ± 10.1 pA, n = 14; with fluctuations: 127.1 pA ± 16.6 pA, n = 14; Figure 5C; Table 2).

**Figure 5:**
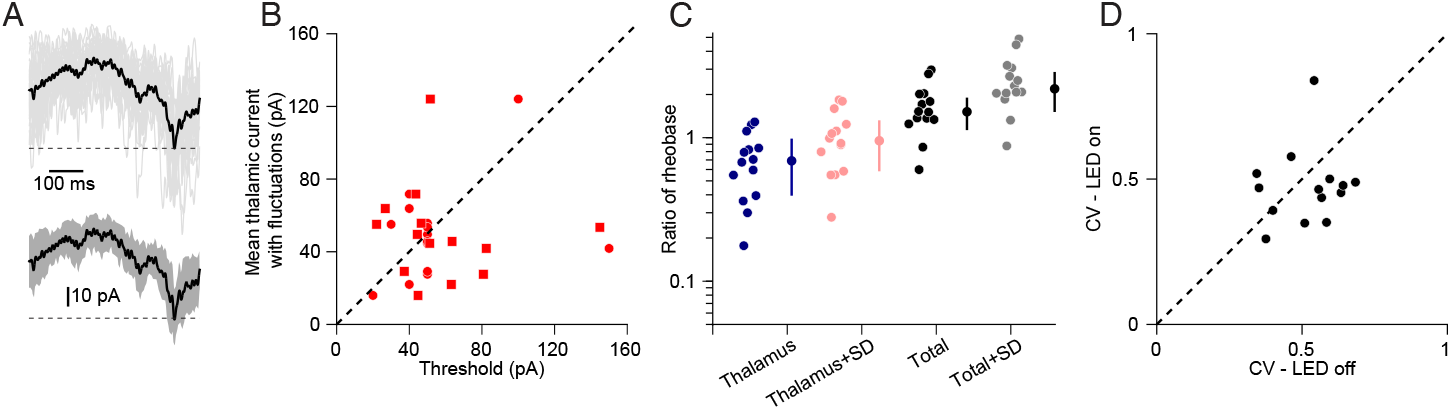
Trial-to-trial fluctuations in thalamic drive are necessary to reach threshold. (A) Top, example cycle-averaged mean response (black) measured from all individual cycle-averaged trials (gray). The dashed line represents the neurons measured statistical threshold. Bottom, same mean response as top example, but with shaded standard deviation (gray). (B) Scatterplot comparing the measured thalamic current during cortical inactivation, with the addition of the standard deviation, to the measured statistical current threshold (filled circles) and fitted current threshold (open squares). (C) Individual log contributions to the current threshold for each neuron under all four measured conditions. Offset circles show the geometric mean for each condition. Error bars: 95% confidence intervals. (D) Comparison of the coefficient of variation for individual neurons during control conditions and optogenetic cortical inactivation.

Because fluctuations in the feedforward drive are important for neurons to reach threshold, we compared the cycle-by-cycle visually-evoked thalamic variability to the overall response variability in our recorded neurons. As there are clear differences in response amplitude between thalamic excitation and total excitation (Figures 4A-C), we chose the CV as our metric of variability in order to normalize these response amplitudes across conditions (Cohen-Kashi Malina et al., 2016). Experiments in cat V1 have shown that cortical variability is inherited from thalamic inputs (Sadagopan & Ferster, 2012). In contrast, other evidence suggests that cortical variability, although inherited partially from thalamic inputs, is locally amplified (Cohen-Kashi Malina et al., 2016). If the trial-to-trial variability in cortex is inherited from the thalamus, the ratio between the thalamic and total CV should be close to 1. When comparing the average CV during cortical silencing and with the cortex intact, we found no significant differences (total CV = 0.51 ± 0.06; thalamic CV = 0.46 ± 0.07; p = 0.2958, n = 14, signed-rank = 70, Wilcoxon signed-rank test; Table 2, Figure 5D). These results indicate that cortical fluctuations could emerge from thalamic drive. We can thus conclude, that the overall variance observed in cortical neurons could be to a large extent inherited from the feedforward inputs in mouse V1. Overall, our results show that while the cycle average thalamic input is subthreshold, temporal fluctuations in that drive are sufficient to evoke spiking in cortical neurons.

### Characterizing the effect of background activity on measured thresholds

It is common to measure the current threshold by injecting current steps of different amplitudes (Jack et al., 1988). This is usually performed *in vitro*, however, where network activity is generally quiescent. During our current injections *in vivo*, neurons are receiving barrages of synaptic input. In many cases, neurons fluctuate between brief periods of depolarized membrane potential, referred to as ‘UP’ states, and brief hyperpolarized periods, referred to as ‘DOWN’ states (Steriade et al., 1993; Cowan & Wilson, 1994; Lampl et al., 1999, Figure 6A). These spontaneous background fluctuations are thought to reflect brief, but powerful, periods of sustained excitatory input onto individual neurons (Wilson & Kawaguchi, 1996; Lampl, et al., 1999). They potentially impact our estimate of current threshold by either changing the distance to the voltage threshold or altering the resistance of neurons. To assess this impact we measured the distribution of membrane potential during spontaneous activity and separated our data based on whether or not recorded membrane potentials were above or below the median (Figure 6A, inset and Figure 6B, see Methods). We use the terms ‘UP’ and ‘DOWN’ state for this separation, though not all of our data formed a bimodal distribution (Figure 6A). We then determined the current threshold separately for UP and DOWN states for each cell.

**Figure 6:**
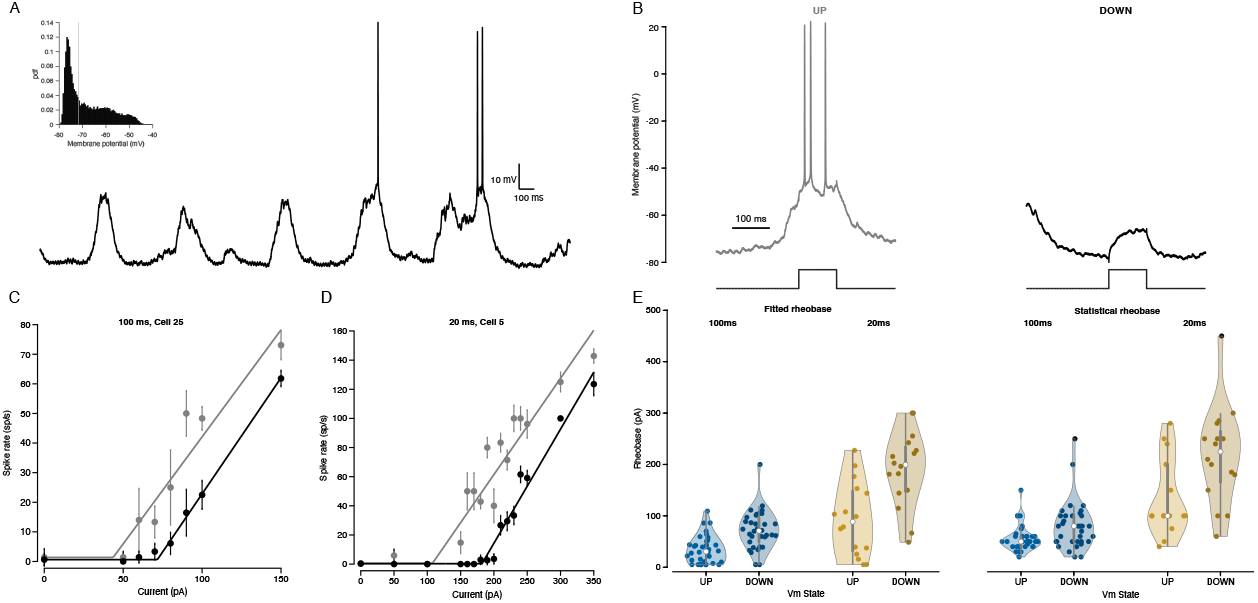
Measured thresholds significantly depend upon background fluctuations in membrane potential state. (A) Example whole-cell trace of spontaneous activity. Under spontaneous conditions during anesthesia, neurons exhibit brief depolarized excursions. Inset, all-points histogram showing the distribution of membrane potential for the example neuron. The gray line represents the median of the distribution. (B) Left, example response to a 100ms pulse of current for the same neuron in (A) during a period of depolarization. Right, example response to current during a period of hyperpolarized membrane potential. (C) Frequency-versus-current curve for an example cell in response to 100ms current pulses. The gray fitted line represents data during membrane potential periods above the median of the membrane potential distribution, while black color line represents data during membrane potential values below the median. Error bars are standard errors. (D) Same as in (C), but for the 20ms pulse duration. (E) Scatterplot comparing thresholds for each neuron measured during membrane potential periods above the median versus below the median for both 100ms pulses (blue) and 20ms pulses (yellow). Circles: measured statistical thresholds. Open squares: fitted thresholds.

We found that, for both the 100ms and 20ms pulse durations, the membrane potential prior to the current injection shifted the threshold for spiking. If the membrane potential prior to a current pulse was in the UP state the current threshold was lower than the DOWN state. This occurred both for the statistical and fitted threshold measurements (see Table 1), consistent with the UP state bringing neurons closer to threshold. Interestingly, we observed the shift in threshold was much higher for the 20ms than the 100ms pulse duration (97.4 and 81.6 pA for 20ms versus 38.1 and 34.1 pA for 100ms; Figure 6E). This is likely due to the timescale of fluctuations responsible for driving neurons closer to threshold: during brief pulses the state of the network does not fluctuate over the course of the input pulse, whereas for the 100ms pulses, the ongoing input can change over the course of the pulse, switching between UP and DOWN states. In sum, we observed that background synaptic drive plays a substantial role in influencing our threshold measurements. Specifically, when neurons are in a depolarized state as a result of ongoing synaptic activity, less current is required to initiate action potentials.

## Discussion

We used intracellular recordings *in vivo* in conjunction with optogenetics to measure the strength of thalamic drive onto visual cortical neurons that receive direct thalamic innervation. We compared this drive to the amount of input necessary to generate action potentials in these neurons. While both thalamic strength and current threshold varied across neurons, the average thalamic drive was consistently below the current threshold to evoke action potentials *in vivo*. We did find, however, considerable trial-to-trial variability in the thalamic drive; the amplitude of this variability, along with the mean drive, is suprathreshold. The variability of the thalamic drive is therefore a critical component to drive cortical action potentials.

Not only is thalamic input variability important for the generation of cortical action potentials, it is also related to the response variability of cortical neurons. Thalamic input, while weaker than the cortical drive, had variability of the same order as the total input variability. Moreover, our finding that input fluctuations in cortical neurons persist when the all recurrent inputs are suppressed suggests that cortical variability could be inherited from the thalamus, similar to what has been reported in cat visual cortex (Sadagopan & Ferster, 2012). Our observations differ, however, from a report in mouse barrel cortex that response variability is higher in the cortex than the thalamus (Cohen-Kashi Malina et al., 2016). While our interpretation differs from this study, our data and analyses are similar. Since the increases in cortical variability observed in barrel cortex may be due to an amplification process we view our reports as differences in degree rather than kind. Cortical variability has often been considered the result of interactions within cortical networks (van Vreeswijk & Sompolinsky, 1996; Hansel & van Vreeswijk, 2012; Hennequin et al., 2018), but our data, as well as those from other groups studying either cat V1 and rodent S1, indicate that variable thalamic drive plays an essential role in generation of cortical responses.

We have emphasized the role of thalamic variability in driving cortical action potentials. An alternative perspective, however, is that the mean drive, while below the current threshold for the majority of neurons, is sufficient to elicit spiking in a small population of thalamo-recipient neurons. Recurrent drive from those cells could then enable other neurons to reach threshold. It is nonetheless the case in this scenario that thalamic input variability will alter the responses of these neurons since it still forms a large component of their overall drive.

Some prior reports of the current threshold differ from our measurements. Studies using sharp electrodes in cat V1 found an average threshold current of 90 ± 20 pA in response to longer current steps (Ahmed et al., 1998). Other studies using sharp electrodes in the motor cortex of adult cats have found even much higher current thresholds, ranging from 600-1800 pA, depending on recorded cell type (Woody & Black-Cleworth, 1973; Baranyi et al., 1993). These values are higher than those we found for the 100 ms pulse duration threshold currents. This may be due to the fact our measurements were performed with whole-cell patch clamp recordings. Indeed, the amount of current needed to evoke spiking activity from sharp recordings is higher than for whole cell recordings (Li et al., 2004). This difference is thought to be because sharp electrodes may introduce membrane leakage resulting in decreased input resistance (Staley et al., 1992; Li et al., 2004).

Background synaptic activity in V1 impacts the response properties of individual neurons (Cowan & Wilson, 1994; Stevens & Zador, 1998; Waters & Helmchen, 2006; Li et al., 2020). Separating our data into states that reflect periods of weak versus strong excitatory input significantly altered the current threshold, particularly for short current injections. While a decline in the current threshold when neurons are depolarized seems expected, changes in input resistance could have altered that relationship (Chance et al., 2002; Mccormick et al., 2003). For example depolarized states have been associated with large conductance changes which could shunt neuronal responses (Hô & Desthexe, 2000). Alternatively it has been shown that depolarization is not necessarily associated with decreases in input resistance (Waters & Helmchen, 2006; Li et al. 2020). Our observations indicate that depolarization, despite resistance changes, reduces current thresholds.

Ongoing noise fluctuations tend to smooth the input-output function of neurons (Anderson et al, 2000b; Hansel & Van Vreeswijk, 2002; Finn et al., 2007). When directly injecting current, however, we observed that the majority of neurons had threshold-linear input-output functions (Fig. 1). Therefore the input fluctuations in the absence of visual stimulation are insufficient to provide the expansive smoothing seen during visually-evoked periods. One reason for the lack of expansive smoothing could be that many of our current injections were suprathreshold, and thus limited the effects of ongoing noise. Another reason, however, is that the variability during visual stimulation is different than during spontaneous activity (Sadagopan & Ferster, 2012; Churchland et al., 2010; Finn et al., 2007; Pattadkal et al., in press). While the variability to smooth the input-output functions has often been ascribed to the cortex (Hennequin et al., 2018, Hansel & van Vreeswijk, 2002), our observations instead indicate that the thalamic drive itself is a source of the noise that generates responses across the visual cortex. Therefore, because the mean thalamic input is subthreshold, the variable afferent drive, found in both somatosensory and visual thalamic inputs, plays an essential role in determining both the mean and variance of cortical responses. This commonality across sensory areas suggests that the root of variability lies in the sensory afferents rather than emerging from cortical networks.

## Materials and Methods

### Animals

All animal procedures were approved by the University of Texas at Austin Institutional Animal Care and Use Committee. Adult mice of both sexes between 5-16 weeks were used for all recordings. For experiments without voltage-clamp measurements, we used C57/BL6J (Jackson Labs #000664). For voltage-clamp inactivation experiments, Parvalbumin(PV)-Cre mice (Scholl et al., 2015) were crossed with a Cre-dependent reporter strain, channelrhodopsin-2 (ChR2)-EYFP strain (Jackson Labs #024109), allowing for selective expression of ChR2 in PV+ interneurons.

### Anesthetized surgical preparation

Surgical procedures for anesthetized mouse preparation are similar to those described previously (Tan et al., 2011). Mice were anesthetized with 1000 mg/kg of urethane and 5 mg/kg of acepromazine for sedation via intraperitoneal injection. Dexamethasone (20 mg/kg) was also administered to prevent brain edema. Temperature was monitored during the entire experiment and body temperature was maintained at 37 degrees celsius using a thermostatically controlled heat lamp. A tracheotomy was performed and the head was placed in a mouse adaptor (Stoelting). A small craniotomy (∼1-3mm) was performed over visual cortex and the dura was removed. A layer of 1.5% agarose in normal saline was placed over the brain to prevent pulsations and to keep the cortex moist. The eyes were kept moist with a thin layer of silicone oil. V1 was located with multiunit extracellular recordings using tungsten electrodes (1MΩ, Micro Probes).

### *In vivo* whole-cell recordings

To obtain recordings, the blind patch method was used. Pipettes were lowered into the cortex and a gigaseal (>1GΩ) was formed onto single neurons. Borosilicate glass capillaries (King Precision Glass) were used to pull electrodes between 5-10 MΩ. Pipettes were filled with a potassium-gluconate based internal solution with the following concentrations (in mM): 135 K-gluconate, 4 NaCl, 0.5 EGTA, 2 MgATP, 10 phosphocreatine disodium, and 10 HEPES. Neurons were recorded 200-800 μm below the cortical surface. Voltage-clamp recordings were restricted to layer 4 of V1, 300-500 μm below the cortical surface. Recordings were made with a MultiClamp 700B patch clamp amplifier (Molecular Devices). For voltage-clamp recordings, neurons were clamped at the reversal potential of inhibition (-75 mv – -80mV) and inward excitatory currents were measured.

### Cortical Silencing

Methods for cortical silencing are the same as those described previously (Barbera et al., 2022). To stimulate ChR2-expressing PV+ interneurons, a 470-nm fiber-coupled LED light was placed above the cortex. During experiments, LED illumination of the cortex was controlled by a TTL signal from a National Instruments board and turned on 250 ms prior to visual stimulation. LED light intensity was between 1 and 1.3 mW.

### Current injection protocol

To measure action potentials in response to current injection, two different pulse durations were used, 100 ms pulses and 20 ms pulses. For each neuron 5-10 trials were performed for each current injection amplitude. Similarly, trials of spontaneous activity were collected. Intertrial intervals were 500 ms long and interstimulus intervals were between 500 – 575 ms to allow for time for the neuron to recover in between current injections. Each trial was 2.5 s long. The coarseness of current steps was tailored for each neuron depending upon measured responsiveness during the recording. Across all neurons, current steps ranged between 10-50 pA current steps, ranging from 0-800 pA current amplitudes.

### Visual stimulation protocol

Current-clamp and voltage-clamp whole-cell recordings were conducted in the binocular zone of V1 (Gordon & Stryker, 1996). Stimuli were generated by a Macintosh computer (Apple) using the Psychophysics Toolbox for MATLAB (MathWorks) and presented on a Sony video monitor (GDM-F520) placed 25 cm from the animal’s eyes. The refresh rate for the monitor was 100 Hz and the spatial resolution was 1024 × 768 pixels. The mean luminance of the monitor was 40 cd/m^2. Drifting gratings were presented to the contralateral eye for 3s at a spatial frequency of 0.04 cycles/deg, temporal frequency of 2 Hz, 100% contrast, and size of 25°. The optimal stimulus orientation was determined by a series of trials of drifting gratings presented in 30° steps. Under voltage-clamp conditions, current measurements were made during visual stimulus presentation of 100% contrast gratings at the previously determined optimal orientation.

### Data analysis

#### Determination of threshold

Action potentials were first segregated from membrane potential data for each current injection in order to determine the number of spikes recorded for each trial. In order to determine the threshold, two separate methods were used. First, trials of spontaneous activity were recorded in order to derive measurements of the spontaneous spike rate. Trials of current injection were made and organized by current amplitude.

The threshold was determined by finding the first current amplitude in which a statistically significant increase between baseline was observed for that current amplitude and subsequent current amplitudes. Statistical significance measurements were Bonferroni corrected for the number of current amplitudes injected to correct for multiple comparisons. In rare cases, due to large variabilities in membrane potential, it was not possible to clearly meet the criterion of consecutive statistically significant current amplitudes, despite observing an increasing spike rate to our current injections in these cells. In these cases, if a statistically significant current was observed and a subsequent current amplitude was not significant, their data were combined to represent an intermediate spike rate. If this dataset was statistically significant, we took the average of the two current amplitudes and set that as the threshold. Second, data were fit to a threshold-linear function, modeled by the following equation:

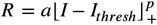

where *R* is the evoked spike rate, *+* is rectification, *p* is the exponent, *I*_*thresh*_ is the fitted threshold current and *a* is a gain factor Data were fitted iteratively over 1000 trials to minimize the mean-squared error between the fitted function using lsqcurvefit in MATLAB.

#### Estimates of input resistance and time constant

In order to derive measurements of intrinsic neuronal properties, we employed a method previously described (Anderson et al., 2000a; Li et al., 2020). We took a subset of our 100ms positive current injections (50 pA – 100 pA) and fit the membrane potential responses to the following equation:

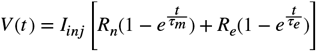

where *V* is the voltage response, *t* is time, *I*_*inj*_ is injected current, *R*_*n*_ is input resistance, *τ*_*m*_ is membrane time constant, *R*_*e*_ is electrode resistance, and *τ*_*e*_ is electrode time constant.

The membrane potential response, in short, can be seen as the sum of these two exponential equations. One corresponds to response to the electrode (e.g., the fast component of the response) while the other corresponds to the response due to the membrane (e.g., the slow component of the response). By fitting the response to this equation, we are able to effectively remove the response to the electrode and derive *τ*_*m*_ and *R*_*n*_.

#### Voltage-clamp current measurements

Inward currents to drifting gratings were measured at the optimal stimulus orientation under conditions when the LED was on and off. Baseline subtraction was performed by measuring the holding current applied in a 250 ms window prior to stimulus onset. The bottom third percentile was taken from the distribution of holding current values in this window on a trial-by-trial basis and this value was subtracted from recordings. Trials were cycle-averaged over a 3s stimulus period (Figure 4A). We used a Fourier transform to calculate the modulation component of the response (F1) and the mean of the response (DC). We considered the sum of these two components to be the total current response to the visual stimulus under each condition (LED on and LED off) separately. The standard deviation was calculated from the time course of the mean response’s standard deviation. We used a Fourier transform to calculate the peak of the standard deviation from the F1 and DC components

#### Threshold across membrane potential states

In order to distinguish threshold current during different membrane conditions, we first median filtered the data using a 10-ms window to remove spikes. Spontaneous membrane potential was computed across the entire dataset to generate a dataset of spontaneous activity. From this, we took the median of the distribution of the spontaneous membrane potential fluctuations and set this value as a voltage threshold for separating our recordings into two separate membrane potential states. We then took an average of the membrane potential values 25ms prior to our current injection and compared it to the median of the spontaneous distribution to categorize the data as either above or below the median. We computed the threshold for these two separate datasets identical to the methods of current threshold analysis described above. In a few cases, spontaneous spike rates were zero during membrane potential down states. In these cases, we took the spontaneous spike rate to be the spontaneous spike rate across our entire distribution of membrane potential values, given that in neurons in which spike rates were nonzero during down states, we found that spike rates were not statistically significantly different from spontaneous rates averaged from the the entire distribution of membrane potential.

## Acknowledgments

We thank Carrie Barr for lab management, animal care, and guidance on surgical procedures. Funding: N.J.P was funded by NIH R01 EY025102 and NS120562; D.H. by ANR-14-NEUC-0001-1 and ANR-17-NEUC-0005; L.J. by 5T32EY021462-13.

## Author Contributions

Conceptualization: L.J, D.H, and N.J.P.

Experimentation: L.J.

Analysis: L.J. and N.J.P.

Writing and editing: L.J., D.H., and N.J.P.

## References

Ahmadian, Y., Rubin, D. B., & Miller, K. D. (2013). Analysis of the Stabilized Supralinear Network. Neural Computation, 25(8), 1994–2037. 10.1162/NECO_a_00472

Ahmed, B., Anderson, J. C., Douglas, R. J., Martin, K. A., & Whitteridge, D. (1998). Estimates of the net excitatory currents evoked by visual stimulation of identified neurons in cat visual cortex. Cerebral Cortex, 8(5), 462–476. 10.1093/cercor/8.5.462

Anderson, J. S., Carandini, M., & Ferster, D. (2000a). Orientation Tuning of Input Conductance, Excitation, and Inhibition in Cat Primary Visual Cortex. Journal of Neurophysiology, 84(2), 909–926. 10.1152/jn.2000.84.2.909

Anderson, J. S., Lampl, I., Gillespie, D. C., & Ferster, D. (2000b). The Contribution of Noise to Contrast Invariance of Orientation Tuning in Cat Visual Cortex. Science, 290(5498), 1968–1972. 10.1126/science.290.5498.1968

Baranyi, A., Szente, M. B., & Woody, C. D. (1993). Electrophysiological characterization of different types of neurons recorded in vivo in the motor cortex of the cat. II. Membrane parameters, action potentials, current-induced voltage responses and electrotonic structures. Journal of Neurophysiology, 69(6), 1865–1879. 10.1152/jn.1993.69.6.1865

Barbera, D., Priebe, N. J., & Glickfeld, L. L. (2022). Feedforward mechanisms of cross-orientation interactions in mouse V1. Neuron, 110(2), 297-311.e4. 10.1016/j.neuron.2021.10.017

Ben-Yishai, R., Bar-Or, R. L., & Sompolinsky, H. (1995). Theory of orientation tuning in visual cortex. Proceedings of the National Academy of Sciences, 92(9), 3844–3848. 10.1073/pnas.92.9.3844

Binzegger, T., Douglas, R. J., & Martin, K. A. C. (2004). A Quantitative Map of the Circuit of Cat Primary Visual Cortex. Journal of Neuroscience, 24(39), 8441–8453. 10.1523/JNEUROSCI.1400-04.2004

Carandini, M., Heeger, D. J., & Movshon, J. A. (1997). Linearity and Normalization in Simple Cells of the Macaque Primary Visual Cortex. Journal of Neuroscience, 17(21), 8621–8644. 10.1523/JNEUROSCI.17-21-08621.1997

Chance, F. S., Abbott, L. F., & Reyes, A. D. (2002). Gain modulation from background synaptic input. Neuron, 35(4), 773–782. 10.1016/s0896-6273(02)00820-6

Churchland, M. M., Yu, B. M., Cunningham, J. P., Sugrue, L. P., Cohen, M. R., Corrado, G. S., Newsome, W. T., Clark, A. M., Hosseini, P., Scott, B. B., Bradley, D. C., Smith, M. A., Kohn, A., Movshon, J. A., Armstrong, K. M., Moore, T., Chang, S. W., Snyder, L. H., Lisberger, S. G., … Shenoy, K. V. (2010). Stimulus onset quenches neural variability: A widespread cortical phenomenon. Nature Neuroscience, 13(3), 369–378. 10.1038/nn.2501

Cohen-Kashi Malina, K., Mohar, B., Rappaport, A. N., & Lampl, I. (2016). Local and thalamic origins of correlated ongoing and sensory-evoked cortical activities. Nature Communications, 7(1), 12740. 10.1038/ncomms12740

Cowan, R. L., & Wilson, C. J. (1994). Spontaneous firing patterns and axonal projections of single corticostriatal neurons in the rat medial agranular cortex. Journal of Neurophysiology, 71(1), 17–32. 10.1152/jn.1994.71.1.17

Douglas, R. J., Koch, C., Mahowald, M., Martin, K. A. C., & Suarez, H. H. (1995). Recurrent Excitation in Neocortical Circuits. Science, 269(5226), 981–985. 10.1126/science.7638624

Ferster, D., Chung, S., & Wheat, H. (1996). Orientation selectivity of thalamic input to simple cells of cat visual cortex. Nature, 380(6571), 249–252. 10.1038/380249a0

Finn, I. M., Priebe, N. J., & Ferster, D. (2007). The emergence of contrast-invariant orientation tuning in simple cells of cat visual cortex. Neuron, 54(1), 137–152. 10.1016/j.neuron.2007.02.029

Gordon, J. A., & Stryker, M. P. (1996). Experience-dependent plasticity of binocular responses in the primary visual cortex of the mouse. Journal of Neuroscience, 16(10), 3274–3286.

Haider, B., Häusser, M., & Carandini, M. (2013). Inhibition dominates sensory responses in the awake cortex. Nature, 493(7430), Article 7430. 10.1038/nature11665

Hansel, D., & Vreeswijk, C. van. (2002). How Noise Contributes to Contrast Invariance of Orientation Tuning in Cat Visual Cortex. Journal of Neuroscience, 22(12), 5118–5128. 10.1523/JNEUROSCI.22-12-05118.2002

Hansel, D., & Van Vreeswijk, C. (2012). The Mechanism of Orientation Selectivity in Primary Visual Cortex without a Functional Map. The Journal of Neuroscience, 32(12), 4049–4064. 10.1523/JNEUROSCI.6284-11.2012

Hennequin, G., Ahmadian, Y., Rubin, D. B., Lengyel, M., & Miller, K. D. (2018). The Dynamical Regime of Sensory Cortex: Stable Dynamics around a Single Stimulus-Tuned Attractor Account for Patterns of Noise Variability. Neuron, 98(4), 846-860.e5. 10.1016/j.neuron.2018.04.017

Hô, N., & Destexhe, A. (2000). Synaptic Background Activity Enhances the Responsiveness of Neocortical Pyramidal Neurons. Journal of Neurophysiology, 84(3), 1488–1496. 10.1152/jn.2000.84.3.1488

Hodgkin, A. L., & Huxley, A. F. (1952). A quantitative description of membrane current and its application to conduction and excitation in nerve. The Journal of Physiology, 117(4), 500–544.

Hodgkin, A. L., & Rushton, W. a. H. (1946). The electrical constants of a crustacean nerve fibre. Proceedings of the Royal Society of London. Series B, Biological Sciences, 133(873), 444– 479. 10.1098/rspb.1946.0024

Hubel, D. H., & Wiesel, T. N. (1962). Receptive fields, binocular interaction and functional architecture in the cat’s visual cortex. The Journal of Physiology, 160(1), 106–154. 10.1113/jphysiol.1962.sp006837

Isaacson, J. S., & Scanziani, M. (2011). How Inhibition Shapes Cortical Activity. Neuron, 72(2), 231–243. 10.1016/j.neuron.2011.09.027

Jack, J. J. B., Noble, D., & Tsien, R. W. (1988). Electric current flow in excitable cells (Reprint). Clarendon Press.

Lampl, I., Reichova, I., & Ferster, D. (1999). Synchronous Membrane Potential Fluctuations in Neurons of the Cat Visual Cortex. Neuron, 22(2), 361–374. 10.1016/S0896-6273(00)81096-X

Li, B., Routh, B. N., Johnston, D., Seidemann, E., & Priebe, N. J. (2020). Voltage-Gated Intrinsic Conductances Shape the Input-Output Relationship of Cortical Neurons in Behaving Primate V1. Neuron, 107(1), 185-196.e4. 10.1016/j.neuron.2020.04.001

Li, L., Li, Y., Zhou, M., Tao, H. W., & Zhang, L. I. (2013). Intracortical multiplication of thalamocortical signals in mouse auditory cortex. Nature Neuroscience, 16(9), 1179–1181. 10.1038/nn.3493

Li, W.-C., Soffe, S. R., & Roberts, A. (2004). A Direct Comparison of Whole Cell Patch and Sharp Electrodes by Simultaneous Recording From Single Spinal Neurons in Frog Tadpoles. Journal of Neurophysiology, 92(1), 380–386. 10.1152/jn.01238.2003

Lien, A. D., & Scanziani, M. (2013). Tuned thalamic excitation is amplified by visual cortical circuits. Nature Neuroscience, 16(9), 1315–1323. 10.1038/nn.3488

Liu, B., Li, Y., Ma, W., Pan, C., Zhang, L. I., & Tao, H. W. (2011). Broad Inhibition Sharpens Orientation Selectivity by Expanding Input Dynamic Range in Mouse Simple Cells. Neuron, 71(3), 542– 554. 10.1016/j.neuron.2011.06.017

Mariño, J., Schummers, J., Lyon, D. C., Schwabe, L., Beck, O., Wiesing, P., Obermayer, K., & Sur, M. (2005). Invariant computations in local cortical networks with balanced excitation and inhibition. Nature Neuroscience, 8(2), 194–201. 10.1038/nn1391

McCormick, D. A., Shu, Y., Hasenstaub, A., Sanchez-Vives, M., Badoual, M., & Bal, T. (2003). Persistent Cortical Activity: Mechanisms of Generation and Effects on Neuronal Excitability. Cerebral Cortex, 13(11), 1219–1231. 10.1093/cercor/bhg104

Miller, K. D., & Troyer, T. W. (2002). Neural Noise Can Explain Expansive, Power-Law Nonlinearities in Neural Response Functions. Journal of Neurophysiology, 87(2), 653–659. 10.1152/jn.00425.2001

Murphy, B. K., & Miller, K. D. (2009). Balanced Amplification: A New Mechanism of Selective Amplification of Neural Activity Patterns. Neuron, 61(4), 635–648. 10.1016/j.neuron.2009.02.005

Pattadkal, J. J., Mato, G., Vreeswijk, C. van, Priebe, N. J., & Hansel, D. (2018). Emergent Orientation Selectivity from Random Networks in Mouse Visual Cortex. Cell Reports, 24(8), 2042-2050.e6. 10.1016/j.celrep.2018.07.054

Pattadkal, J. J., O’Shea, R. T., Hansel, D., Taillefumier, T., Brager, D., & Priebe, N. J. (in press). Synchrony dynamics underlie irregular neocortical spiking (p. 2024.10.15.618398). Neuron.

Peters, A., & Payne, B. R. (1993). Numerical Relationships between Geniculocortical Afferents and Pyramidal Cell Modules in Cat Primary Visual Cortex. Cerebral Cortex, 3(1), 69–78. 10.1093/cercor/3.1.69

Priebe, N. J., & Ferster, D. (2005). Direction Selectivity of Excitation and Inhibition in Simple Cells of the Cat Primary Visual Cortex. Neuron, 45(1), 133–145. 10.1016/j.neuron.2004.12.024

Priebe, N. J., & Ferster, D. (2008). Inhibition, spike threshold, and stimulus selectivity in primary visual cortex. Neuron, 57(4), 482–497. 10.1016/j.neuron.2008.02.005

Priebe, N. J., & Ferster, D. (2012). Mechanisms of neuronal computation in mammalian visual cortex. Neuron, 75(2), 194–208. 10.1016/j.neuron.2012.06.011

Ringach, D. L., Hawken, M. J., & Shapley, R. (1997). Dynamics of orientation tuning in macaque primary visual cortex. Nature, 387(6630), 281–284. 10.1038/387281a0

Sadagopan, S., & Ferster, D. (2012). Feedforward origins of response variability underlying contrast invariant orientation tuning in cat visual cortex. Neuron, 74(5), 911–923. 10.1016/j.neuron.2012.05.007

Scholl, B., Pattadkal, J. J., Dilly, G. A., Priebe, N. J., & Zemelman, B. V. (2015). Local Integration Accounts for Weak Selectivity of Mouse Neocortical Parvalbumin Interneurons. Neuron, 87(2), 424–436. 10.1016/j.neuron.2015.06.030

Shadlen, M. N., & Newsome, W. T. (1998). The Variable Discharge of Cortical Neurons: Implications for Connectivity, Computation, and Information Coding. Journal of Neuroscience, 18(10), 3870– 3896. 10.1523/JNEUROSCI.18-10-03870.1998

Softky, W. (1994). Sub-millisecond coincidence detection in active dendritic trees. Neuroscience, 58(1), 13–41. 10.1016/0306-4522(94)90154-6

Somers, D. C., Nelson, S. B., & Sur, M. (1995). An emergent model of orientation selectivity in cat visual cortical simple cells. Journal of Neuroscience, 15(8), 5448–5465. 10.1523/JNEUROSCI.15-08-05448.1995

Somers, D. C., Todorov, E. V., Siapas, A. G., Toth, L. J., Kim, D. S., & Sur, M. (1998). A local circuit approach to understanding integration of long-range inputs in primary visual cortex. Cerebral Cortex, 8(3), 204–217. 10.1093/cercor/8.3.204

Sompolinsky, H., & Shapley, R. (1997). New perspectives on the mechanisms for orientation selectivity. Current Opinion in Neurobiology, 7(4), 514–522. 10.1016/S0959-4388(97)80031-1

Staley, K. J., Otis, T. S., & Mody, I. (1992). Membrane properties of dentate gyrus granule cells: Comparison of sharp microelectrode and whole-cell recordings. Journal of Neurophysiology, 67(5), 1346–1358. 10.1152/jn.1992.67.5.1346

Steriade, M., Nunez, A., & Amzica, F. (1993). A novel slow (< 1 Hz) oscillation of neocortical neurons in vivo: Depolarizing and hyperpolarizing components. Journal of Neuroscience, 13(8), 3252– 3265. 10.1523/JNEUROSCI.13-08-03252.1993

Stevens, C. F., & Zador, A. M. (1998). Input synchrony and the irregular firing of cortical neurons. Nature Neuroscience, 1(3), 210–217. 10.1038/659

Stratford, K. J., Tarczy-Hornoch, K., Martin, K. A. C., Bannister, N. J., & Jack, J. J. B. (1996). Excitatory synaptic inputs to spiny stellate cells in cat visual cortex. Nature, 382(6588), 258–261. 10.1038/382258a0

Stuart, G., Spruston, N., & Häusser, M. (Eds.). (2016). Dendrites (Third edition). Oxford University Press.

Tan, A. Y. Y., Brown, B. D., Scholl, B., Mohanty, D., & Priebe, N. J. (2011). Orientation Selectivity of Synaptic Input to Neurons in Mouse and Cat Primary Visual Cortex. Journal of Neuroscience, 31(34), 12339–12350. 10.1523/JNEUROSCI.2039-11.2011

Troyer, T. W., & Miller, K. D. (1997). Integrate-and-Fire Neurons Matched to Physiological F-I Curves Yield High Input Sensitivity and Wide Dynamic Range. In J. M. Bower (Ed.), Computational Neuroscience: Trends in Research, 1997 (pp. 197–201). Springer US. 10.1007/978-1-4757-9800-5_32

Van Vreeswijk, C., & Sompolinsky, H. (1996). Chaos in Neuronal Networks with Balanced Excitatory and Inhibitory Activity. Science, 274(5293), 1724–1726. 10.1126/science.274.5293.1724

Waters, J., & Helmchen, F. (2006). Background synaptic activity is sparse in neocortex. The Journal of Neuroscience: The Official Journal of the Society for Neuroscience, 26(32), 8267–8277. 10.1523/JNEUROSCI.2152-06.2006

Wilson, C. J., & Kawaguchi, Y. (1996). The origins of two-state spontaneous membrane potential fluctuations of neostriatal spiny neurons. Journal of Neuroscience, 16(7), 2397–2410. 10.1523/JNEUROSCI.16-07-02397.1996

Woody, C. D., & Black-Cleworth, P. (1973). Differences in excitability of cortical neurons as a function of motor projection in conditioned cats. Journal of Neurophysiology, 36(6), 1104–1116. 10.1152/jn.1973.36.6.1104

